# Temperature-dependence of spontaneous mutation rates

**DOI:** 10.1101/2020.11.03.366807

**Authors:** Ann-Marie Waldvogel, Markus Pfenninger

**Affiliations:** Senckenberg Biodiversity and Climate Research Centre, Georg-Voigt-Str. 14-16, 60325 Frankfurt am Main, Germany; Institute of Zoology, University of Cologne, Zülpicher Str. 47b, 50674 Cologne, Germany; LOEWE Centre for Translational Biodiversity Genomics, Senckenberg Biodiversity and Climate Research Centre, Frankfurt am Main, Germany; Institute for Organismic and Molecular Evolution, Johannes Gutenberg University, Mainz, Germany

## Abstract

Mutation is the source of genetic variation and the fundament of evolution. At the interphase of ecology and evolution, temperature has long been suggested to have a direct impact on realised spontaneous mutation rates. The question is whether mutation rates can be a species-specific constant under variable environmental conditions, such as variation of the ambient temperature. By combining mutation accumulation with whole genome sequencing in a multicellular organism, we provide empirical support to reject this null hypothesis. Instead mutation rates depend on temperature in a U-shaped manner with increasing rates towards both temperature extremes. This relation has important implications for mutation dependent processes in molecular evolution, processes shaping the evolution of mutation rates and even the evolution of biodiversity as such.

---

Mutation has been described as the ‘quantum force’ of biology^1^: pervasive throughout the tree of life, fundamental basis of evolution and notoriously difficult to measure. Evidence for the variation of mutation rates (μ) has been accumulating for a century, with pioneering investigations in *Drosophila* dating back to the 1930s^2,3^ and more recent studies estimating the variability of μ in microorganisms^4–6^, plants^7–9^, invertebrates^10–14^ as well as vertebrates^15–17^. On the intraspecific level, the spontaneous μ was often claimed to be a species-specific constant^18^. The question about which factors directly drive mutational rate variation remains unresolved, particularly in multicellular organisms. Temperature has long been suggested as a major determinant of μ variation. Experiments in the first half of the 20^th^ century (reviewed in Lindgren^19^) reported generally large effects of temperature treatments on phenotypically visible mutations. As reported by Muller^20^ in the first study on this topic, “both the direction of the effect of temperature on the time-rate of mutation, and its approximate magnitude, are the same as in the case of its effect on the time-rate of ordinary chemical reactions”. Interestingly, however, also cold treatments increased the apparent mutation rate in some studies^21,22^. Apart from these early studies, mainly performed even before DNA was identified as the carrier of genetic information, there is surprisingly little empirical evidence if and to what extent temperature modifies the spontaneous rate of mutation on the molecular level. Higher temperature stress had a significant effect on microsatellite μ in *Caenorhabditis elegans*^10^. Studies on *Drosophila melanogaster* suggest that indeed temperature dependent metabolic activities increased the somatic mutation rate as consequence of oxidative stress^23^. Stressful temperature conditions increased μ in experimental evolution lines of the seed beetle *Callosobruchus maculatus*^12^. And the temperature dependent mass specific metabolic rate seemed to influence the interspecific molecular clock rate that depends at least partially on μ^24^.

Despite a century-long research on the potential impact of temperature on mutational variation, we still lack a thorough understanding of mechanisms underlying the temperature-dependence of μ. Empirical evidence for increased μ under stressful temperature conditions is strong, however, the relation between natural, and thus relevant, temperature ranges and the variability of μ on an intraspecific level is still unclear. It was therefore our aim to test the following null hypothesis (H_0_): The spontaneous mutation rate is a species-specific constant and independent of environmental conditions.

To provide a fine-scale resolution of the relation between temperature and μ, we report here directly estimated spontaneous single nucleotide mutation rates in the non-biting midge *Chironomus riparius* under different temperature regimes. *C. riparius* is a fully developed system for multigeneration experiments^25^ with comprehensive genomic resources^26^. A haploid spontaneous single nucleotide μ of 2.1 × 10^−9^ has been previously reported^27^. Moreover, there is indirect evidence for the temperature-dependence of μ among *C. riparius* populations^28^. Here, we performed short-term mutation accumulation experiments at temperatures between 12 °C and 26°C (Fig. 1), representative for the natural range for active development of the species. Temperatures below 12 °C will generally induce developmental pausing for larval overwintering^28^ and temperatures constantly above 26 °C lead to significant negative fitness effects^29^. As expected, the generation time of individuals in our experiments decreased with increasing temperature (Fig. 1) and we also observed a reduction of body size with increasing temperature, a well-known phenotypic response for ectotherms^30^. Whole genome data of the RefPool as ancestral state and 58 single individuals across all mutation accumulation lines (MAL) were passed along the *de novo* mutation calling pipeline established for this species^27^ (and see methods). Across a total number of 5.5 · 10^9^ analysed genomic positions, we identified 106,149 de novo mutations in MAL of all temperatures. Most of these mutations (105,948) were found in five MAL with extraordinary high mutation load (Supplemental Information S1). The occurrence of such so-called mutator lines is a known phenomenon for MAL expereiment^31^ and has also been previously observed in *C. riparius*^32^.

**Figure 1:**
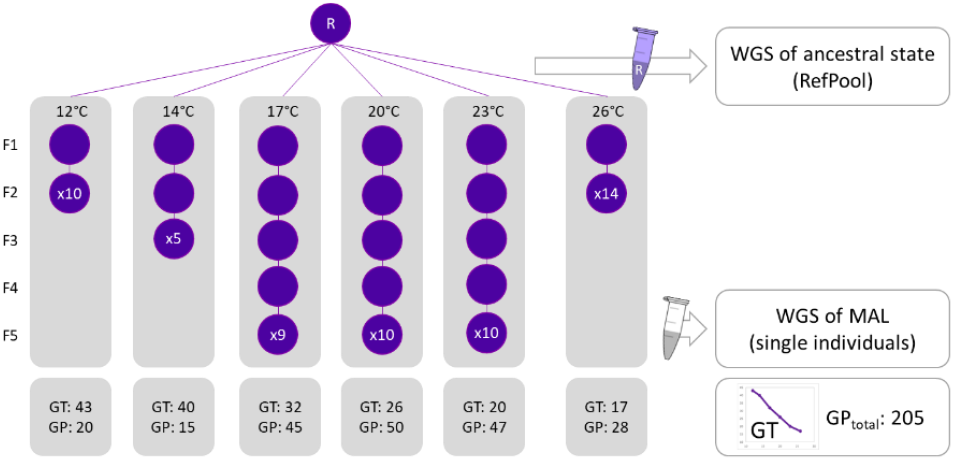
Experimental set up of mutation accumulation at different constant temperatures. The succession of circles indicates how many generations (F1-F5) could be passed under the different temperature conditions. Circles are labelled with the total number of MAL that survived until the respective generation. Mean generation time (GT) is decreasing with increasing temperature as indicated in the detail graph. Number of generational passages (GP = MAL • generation) sums up to a total of 205.

The total rate of spontaneous mutations per haploid genome per generation ranged between 2.77 · 10^−9^ and 1.24 · 10^−8^ across all temperature experiments (Tab. 1). This range covers the previously reported μ for *C. riparius*^27^ and the magnitude is comparable to μ estimates of different other insects, as e.g. *D. melanogaster*^33^ (2.8 · 10^−9^), *Heliconius melpomene*^34^ (2.9 · 10^−9^).

**Table 1:** Overview of results per temperature, excluding mutator lines: **G**enerational **P**assages of successfully sequenced MAL, mean number of **C**allable **S**ites, total number of de novo mutations, number of **S**ingle **N**ucleotide **M**utations, number of **S**ingle **N**ucleotide **I**ndels, mean haploid mutation rates per generation, respectively of total mutations, SNMs and SNIs.

| temperature | GP | mean CS | mutations<br>total | SNMs | SNIs | $\mu_{\text{total}}$ | $\mu_{\text{SNM}}$ | $\mu_{\text{SNI}}$ | transitions | transversions |
| --- | --- | --- | --- | --- | --- | --- | --- | --- | --- | --- |
| 12 | 20 | 8.92E+07 | 27 | 23 | 4 | 7.70E-09 | 6.60E-09 | 1.26E-09 | 17 | 6 |
| 14 | 11 | 7.33E+07 | 6 | 5 | 1 | 4.02E-09 | 3.42E-09 | 9.25E-10 | 3 | 2 |
| 17 | 45 | 1.07E+08 | 26 | 20 | 6 | 2.77E-09 | 2.13E-09 | 6.75E-10 | 8 | 12 |
| 20 | 50 | 1.23E+08 | 52 | 42 | 10 | 4.27E-09 | 3.28E-09 | 8.55E-10 | 18 | 24 |
| 23 | 33 | 9.60E+07 | 40 | 37 | 3 | 6.40E-09 | 5.90E-09 | 5.45E-10 | 26 | 11 |
| 26 | 26 | 7.87E+07 | 50 | 45 | 5 | 1.24E-08 | 1.12E-08 | 1.35E-09 | 28 | 17 |

Temperature had a significant impact on realised spontaneous μ in our MAL experiments. The lowest rate of total mutations was calculated for the 17 °C experiment with 2.77 · 10^−9^ (95% HDI: lower 1.74 · 10^−9^ and upper 3.78 · 10^−9^; Tab. 1 and Supplemental Information S2). From this minimum, μ increased monotonically with the absolute temperature difference in both directions (Fig. 1). μ at the extreme temperatures 12°C (7.70E-09, 95% HDI: lower 4.90 · 10^−9^ and upper 1.06 · 10^8^, Δ5 °C from minimum) and 26°C (1.24E-08, 95% HDI: lower 8.85 · 10^−9^ and upper 1.57 · 10^−8^, Δ 9°C from minimum) was 2.79, respectively 4.54 times higher than the minimum rate (respective posterior probabilities 99.8% and 100%, Tab. 2). In all cases there was predominant to conclusive evidence that the next temperature step in the direction of both extremes induced a higher μ (Tab. 2).

**Table 2:** Selected μ_total_ comparisons of rate ratios between temperatures (cf. Supplemental Information S2 for full table). Posterior probabilities indicate the support of ratios > 1 between compared temperatures.

| Comparison | Rate Ratio | Post. Prob.<br>Rate $> 1$ (%) |
| --- | --- | --- |
| 12°C vs. 17°C | 2.79 | 99.8 |
| 26°C vs. 17°C | 4.54 | 100 |
| 12°C vs. 14°C | 1.92 | 99.9 |
| 14°C vs. 17°C | 1.46 | 53.1 |
| 20°C vs. 17°C | 1.54 | 99.3 |
| 23°C vs. 20°C | 1.49 | 78.7 |
| 26°C vs. 23°C | 1.92 | 97.5 |

When decomposing total μ into SNM (μ_SNM_) and SNIs (μ_SNI_), the distribution of the data in relation to temperature was almost identical for the former (Fig. 2, bottom). Due to the low number of *de novo* SNIs, μ_SNI_ were comparably low (Tab.1, Fig. 2 bottom) and, at first sight, no clear U-shape distribution in relation to temperature appeared. However, fitting second order polynomial regression models to the data found a U-shape relation between temperature and all measured rates, with a respective goodness of fit of r^2^=0.98, r^2^=0.99 and r^2^ = 0.73 (Fig. 2).

**Figure 2:**
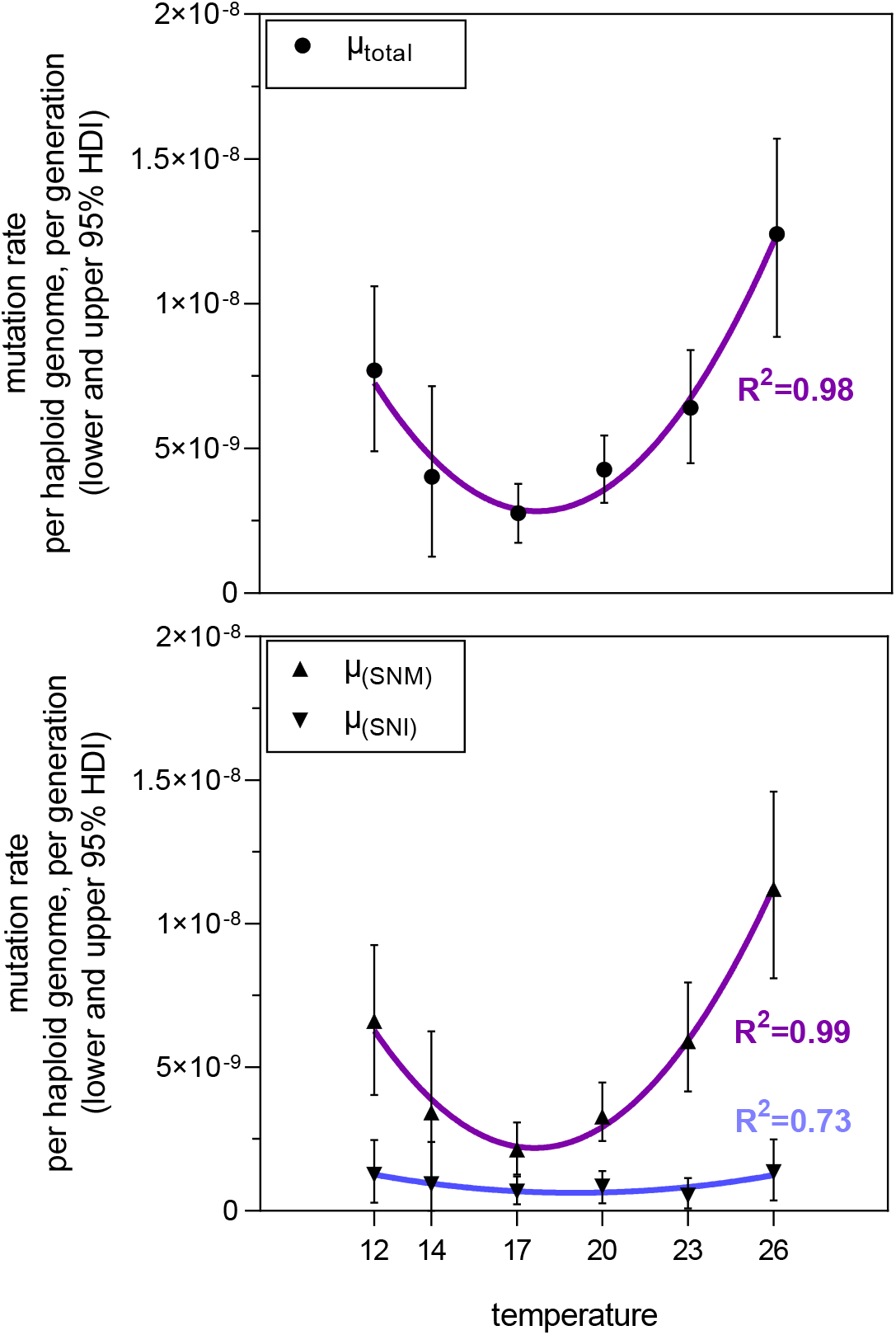
Spontaneous mutation rates per haploid genome per generation in relation to temperature of the MAL experiment. **Top:** Mutation rate of the total sum of de novo mutations. **Bottom:** Separate mutation rates of single nucleotide mutations (SNM) and single nucleotide indels (SNI). Nonlinear fit as second order polynomial regression, R^2^ values are given to describe the goodness of model fit. Corresponding test statistics listed in Supplemental Information S2.

Our results thus deliver unequivocal empirical evidence to reject the H_0_: Single nucleotide mutation rates are not constant within *C. riparius* but depend on temperature with a minimal rate at an optimum temperature. Such a relation was theoretically assumed^35^ but empirical evidence was largely lacking. Before the present study, only Ogur and colleagues^36^ documented the effects of a fine scaled, wider temperature range on spontaneous mutations in *Saccharomyces* species sixty years ago. They used respiration deficiency as phenotypic mutation proxy and found a U-shaped relation, similar to what we found for nucleotide mutations in our whole-genome scale approach in a multicellular organism (Fig. 2). Other studies were restricted either to only partially covering the respective species’ physiological temperature range and/or extreme temperatures that would a priori evoke stressful conditions for the species of choice ^e.g. 10,37^.

There are a few mechanistic hypotheses that could explain the observed pattern^11,35,37^. The metabolic rate hypothesis (MH) suggests that increased metabolic activity leads to the production of free oxygen radicals (ROS) faster than can be effectively eliminated by the organismal antioxidative stress response which act as endogenous mutagens^38^. As ambient temperature determines the metabolic rate of ectotherm organisms, μ should rise with temperature as a consequence of increased oxidative stress, which causes DNA damage and as a result, sometimes erroneous repair^39^. The effect of high temperatures leading to increased mutation rates has been documented for almost a century^2,10,19,20,24^ and recent experiments with *Escherichia coli* underline this^37^. However, it is well established that also low temperatures can induce oxidative stress, even though the mechanisms are not well known^40^. Antioxidative stress response to both high and low temperatures was e.g. observed in *C. riparius*^41^. It was speculated that low temperatures either weakens the systems of ROS elimination, and/or enhances ROS production^38^. In our experiments, the number of successful MAL and their respective amount of generational passages (Fig. 2) reveals the increased stress-level at both of the two extreme temperatures. At 12 and 26 °C, all MAL were lost after the second generation. Whereas the majority of lines reached the fifth generation of mutation accumulation at the intermediate temperatures 17, 20, and 23 °C. Metabolic effects alone could therefore explain the observed pattern.

However, a negative correlation between temperature and μ could also arise from the positive relation between ambient temperature and generation times in the ectotherm *C. riparius*^42^. Whatever mutagens are acting (endogenous ROS production, cosmic rays), their time to induce mutations in the germline should increase with generation time. One could term this the generation length hypothesis (GL). Several studies are lending support for the GL: in primates μ can be predicted via the reproductive longevity^15^ and the mammalian male mutation bias is higher for species with long generation times^43^. There is evidence that this effect does not depend on the absolute number of cell divisions^44^. Also in invertebrates there is significant evidence for a correlation between rates of molecular evolution and generation times^11^. Even though these studies investigated only the relation between temperature independent generation time and μ, their results implied that the temperature-dependent changes in generation time of ectotherms could impact μ. Direct support comes from temperature experiments impacting on life-history and μ in seed beetles^12^. Martin and Palumbi^45^ already concluded their comparative analysis on the relation of body size, metabolic rate, generation time and the molecular clock that our understanding of molecular evolution could be improved when reconsidering “the generation time hypothesis to include physiological effects such as the metabolic rate”.

There is empirical evidence for all of these processes to play a role^35^, leading to potentially complex interactions in each particular case. The here discovered U-shaped relation between μ and temperature could therefore be either due to increased oxidative stress towards physiological extreme temperatures (MH) and/or an interaction with inversely increasing generation times with decreasing temperatures (GL). As generation time and metabolism are intricately linked in ectotherm organisms, additional studies separating these effects are necessary to infer the acting processes.

We additionally disentangled temperature-effects on the mutation spectrum. Excluding mutator lines, we detected 201 de novo mutations during a total of 177 generational passages with 172 SNMs (86 %) and 29 indels (Tab. 1). SNMs divided in 100 transitions and 72 transversions (ratio=1.38, Tab. 1) with a non-random distribution of mutation types among temperatures. Overall, transitions occurred significantly more often than transversions (Mann-Whitney U=53.5, p=0.0015). This pattern provides statistical support for the tendency that we have observed in a previous study with the same species^27^. The transition bias is a well-known pattern of molecular evolution^46^, however, the cause has not yet been resolved. There are two alternative hypotheses relating the bias to either a mutational^47,48^ or selective^49,50^ mechanism. With regard to our experimental design of mutation accumulation, we can neglect selection in favour of “less severe” biochemical amino-acid changes due to transitions, and thus reject the ‘transition-bias due to selection’ hypothesis. More importantly, our data allows a more fine-scaled resolution of the mutation spectrum and we found temperature to have a significant effect on mutation types (F=4.94, p=0.0028), explaining 28 % of the total variation (Fig. 2). Towards the temperature extremes, frequency of all four transition types increased, whereas the effect on transversions was unidirectional with increasing temperatures and mostly due to an increase of A↔T mutations (Fig. 2). This temperature-dependent mutation spectrum provides important evidence that temperature is more than a general stress factor acting on mutational processes, what has been claimed otherwise in a study comparing the effect of temperature stress on spontaneous mutations in nematods^10^. The observed differences in the mutation spectrum suggest that temperature as a bipolar factor is shaping mutational variation through different molecular mechanisms.

Our observation that μ varies significantly in response to environmental temperature has important implications for mutation dependent processes in molecular evolution, processes shaping the evolution of mutation rates and even the evolution of biodiversity as such. Temperature as an abiotic environmental factor is considered to hold a key position at the interphase of ecology and evolution for its influence on organismal physiology^51^, population divergence^52^ and now also mutational variability. The substantial variation of μ in response to ambient temperature, and perhaps also other environmental factors, is especially relevant for our understanding of evolutionary processes under natural conditions. For the multivoltine *C. riparius*, for example, μ in a natural population is expected to vary by a factor of more than four among the generations throughout a year (Fig. 4). This suggests that the mean effective μ over the generations in a year under natural conditions is five times higher than measured in the laboratory^27^. Most interestingly, the effective μ is rather driven by cold temperatures (Fig. 4), with respective consequences for the expected mutational spectrum.

**Figure 3:**
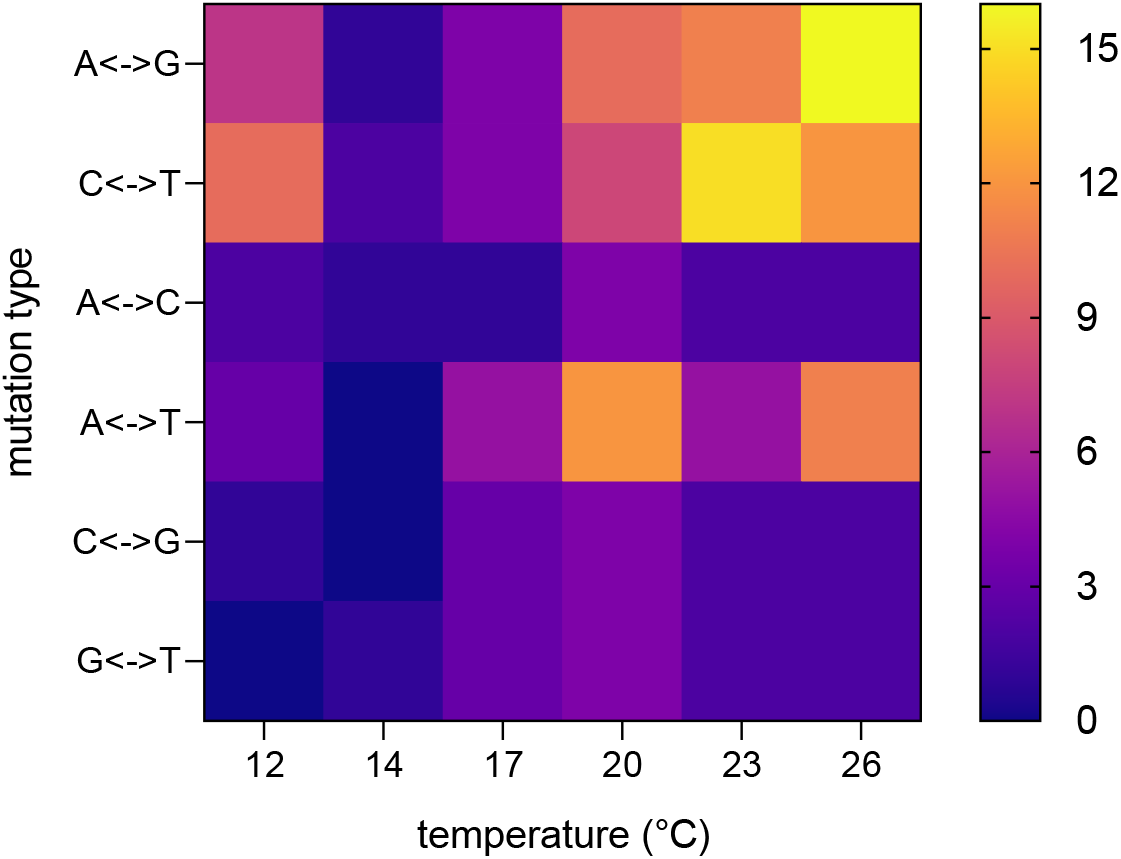
Heatmap of detected SNM types per experimental temperature. Transitions in rows 1 and 2, transversions in rows 3 to 6. Different mutation types explain 43 % (F=7.52, p=0.0002) and temperatures 28 % (F=4.94, p=0.0028) of the total variance.

**Figure 4:**
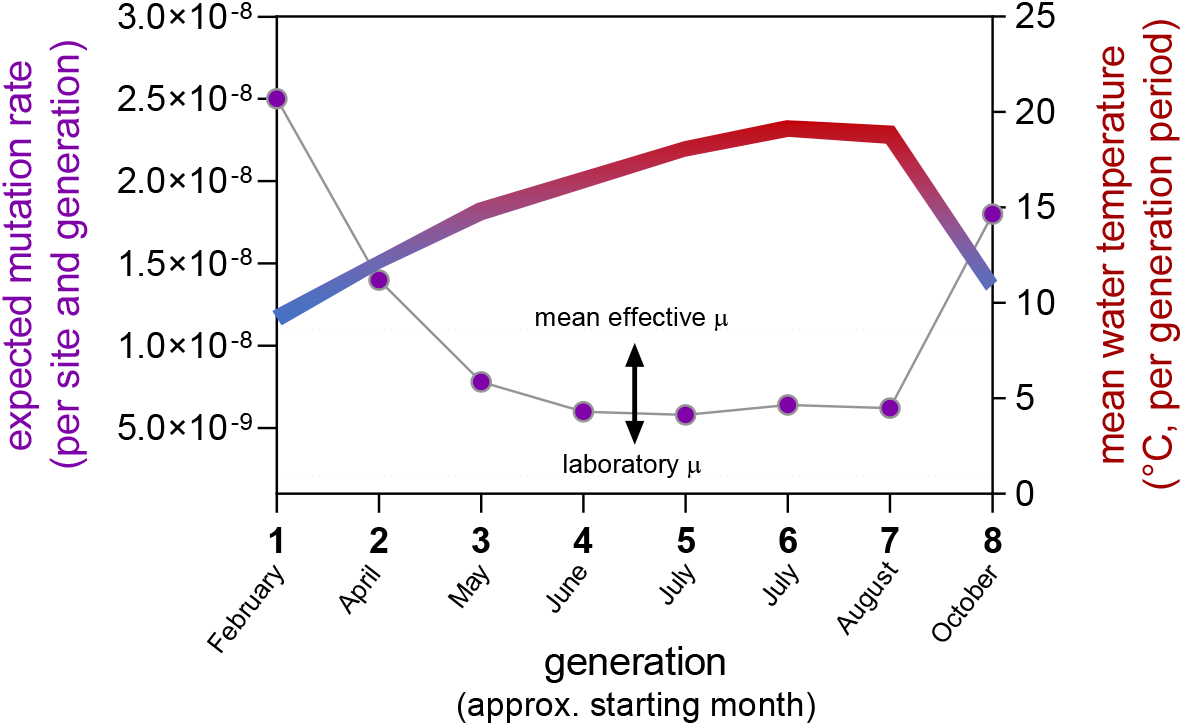
Expected variation of mutation rate calculated per generation for a natural *C. riparius* field population (left y-axis) with regard to the annual variation in water temperature (Hasselbach in Hesse, Germany) averaged for each generation period (right y-axis). Due to the temperature-dependence of generation times, *C. riparius* can pass eight generations per year (x-axis). The mean effective μ is 5.29-fold higher than μ measured under laboratory conditions^27^ (indicated by arrow).

Different other population genetic parameters depend on μ, in particular the estimation of the long-term effective population size (N_e_) from theta_53_. Given the variation of μ with regard to the natural temperature range (Fig. 4), it appears advisable to use a mean effective rate to obtain accurate estimates of N_e_. The same applies for the estimation of divergence times from sequence divergence data, with the additional complication that e.g. changes in N_e_ over time may have additionally influenced the base line μ with regard to the cost-fidelity hypothesis^54^. Last but not least, the evolutionary speed hypothesis (ESH), aiming to explain the latitudinal biodiversity gradient^55^, assumed as one molecular underpinning for faster evolutionary processes in lower latitudes that the higher temperatures in these regions increase μ, leading thus to faster mutation accumulation^28^. If, however, the finding of increased μ towards both temperature extremes is a general phenomenon, it would make this particular aspect of the ESH hypothesis obsolete.

Our scenario of mutation variation to be expected in natural populations also raises the question whether the position of the here observed μ optimum is in itself a species-specific constant or whether this position on the temperature scale can evolve in response to local temperature conditions. Answers to this question will enhance our understanding of the interplay between climate variability and molecular evolution^56^.

## Methods

### Mutation accumulation experiments

The experimental set up for mutation accumulation followed the exact procedure described in Oppold & Pfenninger^27^. In this previous study, we performed the mutation accumulation experiment with the ‘Laufer’ strain of C. riparius under constant conditions at 20 °C. Simultaneously and now presented in this study, we established mutation accumulation lines (MAL) at five additional temperatures: 12 °C, 14 °C, 17 °C, 23 °C and 26°C. Ten MAL were initiated for each temperature condition, with additional backup MAL to compensate for the loss of lines due to lethal inbreeding effects.

### Whole genome resequencing

To establish the genetic baseline of the ancestral state against which to compare de novo mutations, we sequenced the parental individuals as pool of their offspring, hereafter called RefPool. Head capsules of adult midges were pooled for DNA extraction and pooled sequencing on a Illumina HiSeq platform as 150 bp paired-end library to an expected coverage of 60-80X. After mutation accumulation across generations, a single female midge of each MAL was whole genome sequenced on an Illumina HiSeq platform as 150 bp paired-end library to an expected mean coverage of 25X (see Oppold and Pfenninger 2017 for details).

Whole genome sequencing data of RefPools and individuals per MAL was cleaned from adapters and quality trimmed using the automatic wrapper script *autotrim* (available at https://github.com/schellt/autotrim). Genomic data of the 20 °C mutation accumulation experiment (RefPool R1 and 20°C MAL) has already been analysed in our previous study ^27^, however mapped to an older assembly version. To make use of the more complete and less fragmented genome assembly, cleaned reads of RefPools and MAL (including the data published in Oppold & Pfenninger, 2017) were mapped against the latest version of the *C. riparius* genome assembly ^26^ with masked repeat - and TE-regions (see Schmidt et al. 2020 for TE - and repeat-library). We used the Burrows-Wheeler Aligner *bwa mem*^57^ (0.7.17-r1188) for mapping and followed the *GATK* best practices pipeline^58^ to remove duplicates^59^ (picard tools v2.20.8), realignment around indels (insertions and deletions, supported by *GATK* version 4) and recalibration of bases.

### Estimation of spontaneous mutation rates

We define de novo mutations to be single base exchanges (hereafter called single nucleotide mutations, SNM) and single base insertions and deletions (hereafter called single nucleotide indels, SNI) that happen in the germline and are thus inherited to the next generation. After a defined number of generational passages (mutation accumulation) and when compared against an ancestral genotype, de novo mutations can be identified to be novel and unique to a respective MAL. While these criteria are obviously straightforward, their application to whole genome data required a stringent control for base and read quality to avoid false positives as well as false negatives. We therefore combined the statistical approach for genome-wide detection of de novo mutations ^27,33^ with a probabilistic mutation calling approach^60^. Whilst SNIs can only be detected via the statistical approach, the probabilistic mutation calling implemented in the tool *accuMUlate* ^60^ allowed for a more sensitive detection of SNMs lowering the false negative rate. Each set of MAL samples per temperature was compared against its respective RefPool (R1 or R2). For the statistical approach, each MAL sample was analysed as separate bam file, whereas *accuMUlate* requires the input of one overall bam file into which all MAL samples per temperature including the respective RefPool were merged. *accuMUlate* output was further filtered according to the following criteria (cf. Winter et al. 2018 for description of categories): probability of a mutation >=0.9, probability of one mutation >=0.9, probability of correct descendant genotype >=0.9, minimum depth =n_MAL_ · 15X + 15X, maximum depth =n_MAL_ · 47X + 195X, number of mutant allele in RefPool =0, mapping quality difference <=1.96, insert size difference <=1.96, strand bias >=0.05, pair-mapping rate difference >=0.05. For the calculation of mutation rates and to compensate for coverage variation among samples (Supplementary Information S1), we used the separate bam files to more precisely define the number of callable sites for each sample ^details described in 27^ as direct divisor, instead of using the *denominator* of the *accuMUlate* package. Finally, the combination of the two mutation detection approaches gave us the most robust and comprehensive set of de novo mutation candidates, all of which were visually curated in *IGV* (v2.6.3,)^61^, excluding SNMs in the 23 °C mutator lines (see below). A MAL was defined to be a mutator line as soon as we observed a 4-fold increase of its mutation rate when compared against the rates of the remaining MAL of the same temperature.

### Test on temperature dependence of rates

To account for the Poisson distribution of our data (mutation counts), we used a Bayesian framework (R package Bayesian First Aid ^62^) to infer credible intervals (95% highest density intervals of posterior distributions, HDI) and to test differences between rates of different temperatures. To allow for the observation of zero mutations, the model uses Jeffreys prior on lambda. The posterior distribution was sampled with default values (3 chains, 5000 iterations). Posterior probability indicates support of rate ratios. Non-overlapping HDIs of two rates give decisive support for differences between temperatures.

### Estimation of mutational variation for natural population

Daily water temperature data for a *C. riparius* population in a small river (Hasselbach, Hessen, Germany 50.167562°N, 9.083542°E) of a nearby waste water treatment plant was analysed for the years 2009-2018. A population model of the temperature dependence of the generation time was fitted to the averaged data to obtain the number and length of the seasonal generations. The mean water temperature for each generation was then used to infer the expected μ from the second order polynomial function fitted on the relation between experimental temperature and μ (Fig 2, top).

## Data availability

Genomic data generated and analysed for this study are available at European Nucleotide Archive (ENA project number: pending). Detailed lists of de novo mutations and Bayesian estimates can be found in the Supplemental Information linked to this article.

## Acknowledgements

We thank Timm Knautz for his assistance during the experiments and we acknowledge that the “Abwasserverband Freigericht” generously provided the temperature data of the Hasselbach which we used to infer mutation rates in a natural habitat.

## Author contributions

A.-M.W. and M.P. conceived the study, A.M.W. performed the experiments and collected the data, A.-M.W. and M.P. analysed the data and equally contributed to writing the manuscript.

## Supplemental Information

Supplemental Information S1: Detailed lists of de novo mutations per replicate

Supplemental Information S2: Details on statistics of Bayesian estimation and second order polynomial regression

## References

1. Gibson, G. Mutation accumulation of the transcriptome. Nature Genetics vol. 37 458–460 (2005).

2. Timofeeff-Ressovsky, N. W. Qualitativer Vergleich der Mutabilität von Drosophila funebris und D. melanogaster. Z. Indukt. Abstamm. Vererbungsl. 71, 276–280 (1936).

3. Demerec, M. Frequency of spontaneous mutations in certain stocks of Drosophila melanogaster. Genetics 22, 469–78 (1937).

4. Ramiro, R. S., Durão, P., Bank, C. & Gordo, I. Low mutational load and high mutation rate variation in gut commensal bacteria. PLoS Biol. 18, e3000617 (2020).

5. Dillon, M. M., Sung, W., Lynch, M. & Cooper, V. S. Periodic variation of mutation rates in bacterial genomes associated with replication timing. MBio 9, 2020 (2018).

6. Lee, H., Popodi, E., Tang, H. & Foster, P. L. Rate and molecular spectrum of spontaneous mutations in the bacterium Escherichia coli as determined by whole-genome sequencing. Proc. Natl. Acad. Sci. U. S. A. 109, E2774–E2783 (2012).

7. Wang, L. et al. The architecture of intra-organism mutation rate variation in plants. PLOS Biol. 17, e3000191 (2019).

8. Sutherland, W. J. & Watkinson, A. R. Somatic mutation: Do plants evolve differently? Nature 320, 305 (1986).

9. Krasovec, M., Rickaby, R. E. M. & Filatov, D. A. Evolution of mutation rate in astronomically large phytoplankton populations. Genome Biol. Evol. 12, 1051–1059 (2020).

10. Matsuba, C., Ostrow, D. G., Salomon, M. P., Tolani, A. & Baer, C. F. Temperature, stress and spontaneous mutation in Caenorhabditis briggsae and Caenorhabditis elegans. Biol. Lett. 9, (2013).

11. Thomas, J. A., Welch, J. J., Lanfear, R. & Bromham, L. A generation time effect on the rate of molecular evolution in invertebrates. Mol. Biol. Evol. 27, 1173–1180 (2010).

12. Berger, D., Stångberg, J., Grieshop, K., Martinossi-Allibert, I. & Arnqvist, G. Temperature effects on life-history trade-offs, germline maintenance and mutation rate under simulated climate warming. Proc. R. Soc. B Biol. Sci. 284, (2017).

13. Flynn, J. M., Chain, F. J. J., Schoen, D. J. & Cristescu, M. E. Spontaneous mutation accumulation in Daphnia pulex in selection-free vs. competitive environments. Mol. Biol. Evol. 34, 160–173 (2017).

14. Pfenninger, M., Binde Doria, H., Nickel, J., Thielsch, A. & Schwenk, K. Spontaneous rate of clonal mutations in Daphnia galeata. bioRxiv 2020.08.31.275495 (2020) doi:10.1101/2020.08.31.275495.

15. Thomas, G. W. C. et al. Reproductive longevity predicts mutation rates in primates. Curr. Biol. 28, 3193-3197.e5 (2018).

16. Hodgkinson, A. & Eyre-Walker, A. Variation in the mutation rate across mammalian genomes. Nat. Rev. Genet. 12, 756–766 (2011).

17. Carlson, J. et al. Extremely rare variants reveal patterns of germline mutation rate heterogeneity in humans. Nat. Commun. 9, 1–13 (2018).

18. Lynch, M. Evolution of the mutation rate. Trends Genet. 26, 345–352 (2010).

19. Lindgren, D. The temperature influence on the spontaneous mutation rate. Hereditas 70, 165–178 (1972).

20. Muller, H. J. The Measurement of gene mutation rate in Drosophila, its high variability, and its dependence upon temperature. Genetics 13, 279–357 (1928).

21. Birkina, B. N. The effect of low temperature on the mutation process in Drosophila melanogaster. Biol.Zh. Moscow 7, 653–60000 (1938).

22. Kerkis, J. The effect of low temperature on the mutation frequency in D. melanogaster with consideration about the cause of mutation in nature. Drosoph. Inform. Serv. 15, 25 (1941).

23. Garcia, A. M. et al. Age- and temperature-dependent somatic mutation accumulation in Drosophila melanogaster. PLoS Genet. 6, 27 (2010).

24. Gillooly, J. F., Allen, A. P., West, G. B. & Brown, J. H. The rate of DNA evolution: Effects of body size and temperature on the molecular clock. Proc. Natl. Acad. Sci. U. S. A. 102, 140–145 (2005).

25. Foucault, Q., Wieser, A., Waldvogel, A.-M. & Pfenninger, M. Establishing laboratory cultures and performing ecological and evolutionary experiments with the emerging model species Chironomus riparius. J. Appl. Entomol. 143, (2019).

26. Schmidt, H. et al. A high-quality genome assembly from short and long reads for the non-biting midge Chironomus riparius (Diptera). G3-Genes Genomes Genet. 10, 1151–1157 (2020).

27. Oppold, A. M. & Pfenninger, M. Direct estimation of the spontaneous mutation rate by short-term mutation accumulation lines in Chironomus riparius. Evol. Lett. 1, 86–92 (2017).

28. Oppold, A.-M. et al. Support for the evolutionary speed hypothesis from intraspecific population genetic data in the non-biting midge Chironomus riparius. Proceedings. Biol. Sci. 283, (2016).

29. Nemec, S., Patel, S., Nowak, C. & Pfenninger, M. Evolutionary determinants of population differences in population growth rate x habitat temperature interactions in Chironomus riparius. Oecologia 172, 585–594 (2013).

30. Atkinson, D. Temperature and organism size - a biological law for ectotherms? in Advances in Ecological Research vol. 25 1–58 (Academic Press, 1994).

31. Haag-Liautard, C. et al. Direct estimation of the mitochondrial DNA mutation rate inb Drosophila melanogaster. PLoS Biol. 6, 1706–1714 (2008).

32. Binde Doria, H., Waldvogel, A.-M. & Pfenninger, M. Measuring mutagenicity in ecotoxicology: A case study of Cd exposure in Chironomus riparius. bioRxiv (2020) doi:10.1101/2020.11.02.365379.

33. Keightley, P. D., Ness, R. W., Halligan, D. L. & Haddrill, P. R. Estimation of the spontaneous mutation rate per nucleotide site in a Drosophila melanogaster full-sib family. Genetics 196, 313–320 (2014).

34. Keightley, P. D. et al. Estimation of the spontaneous mutation rate in Heliconius melpomene. Mol. Biol. Evol. 32, 239–243 (2015).

35. Baer, C. F., Miyamoto, M. M. & Denver, D. R. Mutation rate variation in multicellular eukaryotes: causes and consequences. Nat. Rev. Genet. 8, 619–631 (2007).

36. Ogur, M., Ogur, S. & St. John, R. Temperature dependence of the spontaneous mutation rate to repsiration deficiency in Saccharomyces. Genetics 45, (1960).

37. Chu, X. L. et al. Temperature responses of mutation rate and mutational spectrum in an Escherichia coli strain and the correlation with metabolic rate. BMC Evol. Biol. 18, 1–8 (2018).

38. Lushchak, V. I. Environmentally induced oxidative stress in aquatic animals. Aquatic Toxicology vol. 101 13–30 (2011).

39. Barzilai, A. & Yamamoto, K. I. DNA damage responses to oxidative stress. DNA Repair (Amst). 3, 1109–1115 (2004).

40. Lalouette, L., Williams, C. M., Hervant, F., Sinclair, B. J. & Renault, D. Metabolic rate and oxidative stress in insects exposed to low temperature thermal fluctuations. Comp. Biochem. Physiol. - A Mol. Integr. Physiol. 158, 229–234 (2011).

41. Park, K. & Kwak, I. S. The effect of temperature gradients on endocrine signaling and antioxidant gene expression during Chironomus riparius development. Sci. Total Environ. 470–471, 1003–1011 (2014).

42. Oppold, A.-M. et al. Support for the evolutionary speed hypothesis from intraspecific population genetic data in the non-biting midge Chironomus riparius. Proc. R. Soc. B Biol. Sci. 283, (2016).

43. Goetting-Minesky, M. P. & Makova, K. D. Mammalian male mutation bias: Impacts of generation time and regional variation in substitution rates. J. Mol. Evol. 63, 537–544 (2006).

44. Wu, F. L. et al. A comparison of humans and baboons suggests germline mutation rates do not track cell divisions. PLoS Biol. 18, e3000838 (2020).

45. Martin, A. P. & Palumbi, S. R. Body size, metabolic rate, generation time, and the molecular clock. Proc. Natl. Acad. Sci. U. S. A. 90, 4087–4091 (1993).

46. Stoltzfus, A. & Norris, R. W. On the Causes of Evolutionary Transition:Transversion Bias. Mol. Biol. Evol. 33, 595–602 (2016).

47. Vogel, F. & Kopun, M. Higher frequencies of transitions among point mutations. J. Mol. Evol. 9, 159–180 (1977).

48. Gojobori, T., Li, W. H. & Graur, D. Patterns of nucleotide substitution in pseudogenes and functional genes. J. Mol. Evol. 18, 360–369 (1982).

49. Rosenberg, M. S., Subramanian, S. & Kumar, S. Patterns of transitional mutation biases within and among mammalian genomes. Mol. Biol. Evol. 20, 988–993 (2003).

50. Wakeley, J. The excess of transitions among nucleotide substitutions: New methods of estimating transition bias underscore its significance. Trends in Ecology and Evolution vol. 11 158–162 (1996).

51. Clarke, A. & Fraser, K. P. P. Why does metabolism scale with temperature? Funct. Ecol. 18, 243–251 (2004).

52. Waldvogel, A. M. et al. The genomic footprint of climate adaptation in Chironomus riparius. Mol. Ecol. 27, 1439–1456 (2018).

53. Charlesworth, B. Effective population size and patterns of molecular evolution and variation. Nat. Rev. Genet. 10, 195–205 (2009).

54. Sung, W., Ackerman, M. S., Miller, S. F., Doak, T. G. & Lynch, M. Drift-barrier hypothesis and mutation rate evolution. Proc. Natl. Acad. Sci. U. S. A. 109, 18488–18492 (2012).

55. Rhode, K. Latitudinal Gradients in Species Diversity: The Search for the Primary Cause. Oikos 65, 514–527 (1992).

56. Pauls, S. U., Nowak, C., Bálint, M. & Pfenninger, M. The impact of global climate change on genetic diversity within populations and species. Molecular Ecology vol. 22 925–946 (2013).

57. Li, H. & Durbin, R. Fast and accurate short read alignment with Burrows-Wheeler transform. Bioinformatics 25, 1754–1760 (2009).

58. McKenna, A. et al. The Genome Analysis Toolkit: A MapReduce framework for analyzing next-generation DNA sequencing data. Genome Res. 20, 1297–1303 (2010).

59. Institute, B. Picard Toolkit. http://broadinstitute.github.io/picard/ (2019).

60. Winter, D. J. et al. accuMUlate: a mutation caller designed for mutation accumulation experiments. Bioinformatics 34, 2659–2660 (2018).

61. Thorvaldsdottir, H., Robinson, J. T. & Mesirov, J. P. Integrative Genomics Viewer (IGV): high-performance genomics data visualization and exploration. Brief. Bioinform. 14, 178–192 (2013).

62. Bååth, R. Bayesian First Aid : A package that implements Bayesian alternatives to the classical *. test functions in R. Proc. UseR 2014 33, 2 (2014).

